# *Fonsecaea pedrosoi* conidia and hyphae activate neutrophils distinctly: Requirement of TLR-2 and TLR-4 in neutrophil effector functions

**DOI:** 10.1101/2020.01.06.895706

**Authors:** Leandro Carvalho Dantas Breda, Cristiane Naffah de Souza Breda, José Roberto Fogaça de Almeida, Larissa Neves Monteiro Paulo, Grasielle Pereira Jannuzzi, Isabela de Godoy Menezes, Renata Chaves Albuquerque, Niels Olsen Saraiva Câmara, Karen Spadari Ferreira, Sandro Rogério de Almeida

## Abstract

Chromoblastomycosis is a chronic and progressive subcutaneous mycosis caused mainly by the fungus *Fonsecaea pedrosoi*. The infection is characterized by erythaematous papules and the histological sections demonstrating an external layer of fibrous tissue and an internal layer of thick granulomatous inflammatory tissue containing mainly macrophages and neutrophils. Several groups have been studying the roles of the innate and adaptive immune systems in *F. pedrosoi* infection; however, few studies have focused on the role of neutrophils in this infection. In the current study, we verified the importance of murine neutrophils in the killing of *F. pedrosoi* conidia and hyphae. We demonstrate that phagocytosis and reactive oxygen species during infection with conidia are TLR-2 and TLR-4-dependent and are essentials for conidial killing. Meanwhile, hyphal killing occurs by NETs formation, in a TLR-2, TLR-4 and ROS-independent manner. In vivo experiments showed that TLR-2 and TLR-4 are also important in Chromoblastomycosis infection. TLR-2KO and TLR-4KO animals had lower levels of MIP-2 and KC chemokines and impaired neutrophil migration to the infected site. These animals also had higher fungal loads during infection with *F. pedrosoi* conidia, confirming that TLR-2 and TLR-4 are essential receptors for *F. pedrosoi* recognition and immune system activation. Therefore, this study demonstrated for the first time that neutrophils activation during *F. pedrosoi* is conidial or hyphal-specific, with TLR-2 and TLR-4 being essential during conidial infection but unnecessary for hyphal killing by neutrophils

**Author Summary:** Chromoblastomycosis (CBM) is a chronic and progressive subcutaneous mycosis that affects mainly low-income individuals, such as farm workers. CBM have been diagnosed all over the world, but the majority of cases were diagnosed in tropical and subtropical climate countries. The treatment is difficult and involves the combination of antifungal prescriptions, cryo/heat-therapy and, in some cases, surgery to remove all the infected tissue. The treatment is long (at least 6 months) and expensive, leading to a high rate of treatment dropout and disease relapse. However, the understanding of pathogen-host interaction is far from being elucitaded. Our understanding is that this pathogen-host interaction in CBM needs to be uncovered so a different and more effective treatment could be proposed to help those patients that are struggling against this chronic infection. Therefore, our study shed a light in neutrophils and innate immune response against this fungal infection, showing the neutrophils capacity to kill *Fonsecaea pedrosoi* conidia and hifa, showing the importance of toll-like receptors 2 and 4 in the neutrophil fungicidal capacity against *F. pedrosoi* conidia but not hyphae. Therefore this work help to the better understanding of how our organism fights back in the CBM infection.

## Introduction

Chromoblastomycosis (CBM) is a chronic, progressive subcutaneous mycosis caused by different fungal species of the *Herpotrichiellaceae* family, such as *Phialophora verrucosa*, *Cladophialophora carrionii, Rhinocladiella aquaspersa, Exophiala spinifera, Aureobasidium pullulans, Chaetomium funicola, Fonsecaea. monophora, Fonsecaea nubica* and *Fonsecaea pugnacious,* but mainly by *Fonsecaea pedrosoi* (1, 2). The disease has been diagnosed on all 5 continents but it is mainly founded in tropical and subtropical countries (3) such as Brazil (4, 5), Mexico (6), China (7) and Madagascar (8). It affects mostly farm workers, because the natural habitat of this fungi is in the soil and decaying plants (9). The treatment is difficult and involves the combination of antifungal prescriptions (10), cryo/heat-therapy (11) and, in some cases, surgery to remove all the infected tissue (12). CBM is one of the most difficult deep mycosis to treat and has low rates of cure (13, 14). The treatment is long and expensive, and because the disease affects mainly low-income individuals, there is a high rate of treatment dropout, leading to a high rate of disease relapse (14). Therefore, a better understanding of the pathogen-host interaction is needed to improve the treatment of CBM, to increase the rate of successful treatment rate and decrease the time and cost of treatment. It is well established in the literature that T-cells and IFN-γ are important for disease control (15–17), but little is known about the innate immune response in CBM. *De Souza’s* lab showed that *F. pedrosoi* conidia ingested by resident macrophages are able to grow into hyphae, leading to macrophages death (18). However, IFN-γ pre-activated macrophages had a fungistatic activity, decreasing hyphal growth and remaining alive (19). Neutrophils are another important innate immune cells during an infection process. They are the most abundant leukocyte in the bloodstream, and the first cells to migrate towards the infection site (20).

Neutrophils are directly responsible for pathogen killing, mainly through three different effector functions: 1) Phagocytosis, 2) Degranulation, and 3) Neutrophil Extracellular Trap (NET) release. These cells can also indirectly control an infection by secreting IL-17 that attracts Th17 lymphocytes, which are an important cell population for fungal infection control (21, 22). Neutrophils can also modulate macrophage phenotypes, helping the immune system against the infection (23). However, although neutrophils activation is usually associated with pathogen containment and elimination, overactivation may be harmful to the host (24, 25), so a tight regulatory system for neutrophil activation is important (26). Although neutrophils are known to be important in several fungal infections such as *Candida albicans* (27), *Aspergillus fumigatus.* (28), *Cryptococcus neoformans* (29), *Paracoccidioides brasiliensis* (30) and *Sporothrix schenkii* (31), few studies have focused on the neutrophils response during CBM infection. However, besides been founded in CBM skin lesions, the significance of neutrophils in helping the immune system avoid fungal spread and promote fungal killing in CBM is unknown (32, 33). Rozental and colleagues demonstrated that neutrophils phagocytise conidia and produce ROS to kill the ingested conidia (34). However, which receptors are responsible for neutrophil activation and whether these cells are able to recognize and kill *F. pedrosoi* hyphal structures is still unknown. In this study, we demonstrate for the first time that conidial killing by neutrophils is TLR-2 and TLR-4-dependent. We also showed that neutrophils’ hyphal killing occurs by NETs release in a TLR-2, TLR-4 and ROS-independent manner. Taken together, our findings help to better understand the neutrophils response in the control of CBM disease, caused by conidial or hyphal infection.

## Materials and Methods

### Research Ethics Board Approval

The protocol for animals studies was approved by the ethics committee (“Comissão de Ética no Uso de Animais da Faculdade de Ciências Farmacêuticas da Universidade de São Paulo”) under protocol number 474. The study was conducted in accordance with the “Conselho Nacional de Controle de Experimentação Animal” (CONCEA) and the “Sociedade Brasileira de Ciências em Animais de Laboratório” (SBCAL) guidelines.

### Fungal strain and growing conditions

The *F. pedrosoi* strain (CBS 271.37) was cultivated on Sabouraud agar at 30°C until the inoculum preparation. The fungi were transferred from the Sabouraud agar tube to 150 mL of Potato Dextrose broth (Difco, BD) and grown for 5 days at 30°C with shaking. After the growth period, the inoculum was filtered through a 40 μM cell strainer. The conidial particles were obtained from the flow-through solution, while the hyphae were retained in the cell-strainer. Conidia-enriched samples were centrifuged for 5 minutes in 300 x *g* to collect the remains of small hyphae and large conidia. The supernatant was collected and centrifuged for 10 minutes at 9000 x *g* and then resuspended in 1x PBS. The concentration of conidia and hyphae was determined by Neubauer chamber counting.

### Mouse bone-marrow neutrophil enrichment

Wild-type, TLR-2KO and TLR-4KO C57BL/6 animals at 8-12 weeks of life were used in this study. The animals were euthanized with an overdose of anaesthetics according to animal ethics committee approval. Femurs and tibias were taken, and the bone marrow was collected using fetal bovine serum-free RPMI medium. The cells were passed through a cell strainer to retain the small debris and clots. The cells were washed once in 1x PBS and neutrophil enrichment was performed using a Ficoll density layer (1119 and 1077 density) or by positive selection with anti-Ly6G magnetics beads (according to the manufacturer’s instructions – Miltenyi®). After neutrophils enrichment, the cells were counted using a Neubauer chamber with trypan blue staining to calculate cell viability. Sample purity was analysed by flow cytometry (anti-CD11b and anti-Ly6G) or the cytospin technique. The viability and purity of the cells used in this work exceeded 95% and 85%, respectively

### Neutrophil fungicidal assay

Purified neutrophils (1×10^5^) were infected with *F. pedrosoi* conidia (multiplicity of infection (MOI) 1:2) or hyphae (MOI 1:1) for 2 hours at 37° C under homogenization. Different MOIs were previously tested with similar results. Therefore, MOIs of 1:2 and 1:1 were chosen for conidia and hyphae experiments, respectively. As a control, conidia or hyphae were incubated under the same conditions without neutrophils. After incubation, an aliquot was taken and diluted in distilled water to induce neutrophil lysis, without harming the fungi. The fungi were seeded onto Sabouraud agar and incubated for 5 days at 37° C for colony-forming unit (CFU) counting. The CFUs of the control groups (fungi without neutrophils) were set as 100% CFU (100% survival). To confirm whether the conidial and hyphal killing was due phagocytosis or NETs release, we first incubated the neutrophils with cytochalasin D (5 μg/mL or DMSO as vehicle) or DNase (25 U/mL) for 15 minutes. The neutrophils were then lysed with distilled water and the fungi were seeded onto Sabouraud agar. To demonstrated that ROS production during the phagocytosis was essential to conidia killing a survival assay was performed as described above in the presence of a range of concentration (0 - 20 μM) of diphenyleneiodonium (DPI), an inhibitor of NADPH-oxidase.

### Phagocytosis and neutrophil extracellular trap (NET) assay

Purified neutrophils from WT, TLR-2KO and TLR-4KO (1.5×10^5^) mice were seeded onto round coverslips previously treated with poly-L-lysine (Sigma-Aldrich®) and placed at the bottom of 24 well plates. After, conidia (MOI 1:2) or hyphae (MOI 5:1) were added and the plates were quickly centrifuged to increase the cell adhesion on the coverslip. After 2 hrs, the supernatant was removed and the cells were fixed with 4% paraformaldehyde (PFA) for 15 minutes. After washing with PBS, the cells were permeabilized with 0,1% PBS-T for 15 minutes and the DNA was stained with Sytox Green (4 μM) for 30 minutes. After washing, the coverslip was placed over a slide with 5 μL of Vecta-Shield® and sealed with nail polish. The slides were kept in the dark at 4° C until analysis by immunofluorescence microscopy. For NETs detection, neutrophils were incubated with hyphae (MOI 5:1) for 1 hour. After, cells were washed and fixed with 4% PFA for 15 min. After washing, Sytox Green (4 μM) was added for 30 minutes and the slides were mounted with Vecta-Shield. The phagocytic index was calculated using the following equation: (number of conidia inside the cells x 100)/ total number of neutrophils. The phagocytic index calculated in WT animals was set as 1 and the TLR-2KO and TLR-4KO phagocytic indexes were then compared to the WT. For the NETs quantification assay, 0.5x x10^5^ neutrophils (WT, TLR-2KO or TLR-4KO) in RPMI medium were seeded into 96 well-plates with Sytox Green (4 μM). The cells were stimulated with conidia (MOI 1:2), hyphae (MOI 5:1) or medium only (negative control) and kept at 37°C with 5% (v/v) CO_2_ incubator and Sytox Green fluorescence intensities were detected by a SpectraMax M2 fluorescence microplate reader (Molecular Devices) every 30 min for 180 min. The NETotic ratio was calculated based on the value of NET formation in unstimulated neutrophils at each specific timepoint (ratio of 1).

### Reactive Oxygen Species (ROS) detection assay

ROS production (O_2_^−^, H_2_O_2_, HOCl) is usually associated with phagocyted pathogen killing. A luminol-enhanced chemiluminescence assay was used to measure the ROS production during conidia and hyphae infection. Briefly, 1×10^5^ neutrophils were seeded into a white 96-well plate (Costar 3917) in the presence of the luminol reagent (1 mmol/L; Sigma-Aldrich). Conidia (MOI 1:2) or hyphae (MOI 5:1) were added and the chemiluminescence was detected with a microplate luminometer reader (EG&G Berthold LB96V, Bad Wildbad, Germany) every 2 minutes for 90 minutes. To measure the ROS production in resting neutrophils, the cells were incubated without any stimuli. The area under curve was calculated to measure the total ROS production after the 90 min stimulation period. To compare the ROS production between WT, TLR-2KO and TLR-4KO, an unstimulated neutrophils sample of each group was set as a ratio of 1. The total ROS production of each group after conidia or hyphae stimulation was compared to its unstimulated sample. To analyze whether hyphae blocked ROS production, neutrophils were stimulated with phorbol 12-myristate 13-acetate (PMA; 100 nM), a well-known NADPH-oxidase agonist, in the presence of hyphae or heat-killed hyphae (HK-hyphae). HK-hyphae were obtained by heating hyphae for 120 minutes in a 90° C in a dry bath block. After heat killing, an aliquot was seeded onto Sabouraud agar plates to confirm that the hyphae were dead (data not shown).

### Chemiotaxis assay

To verify whether the TLR-2 and TLR-4 were essential for neutrophil migration to the infection site, WT, TLR-2KO and TLR-4KO animals were intraperitoneally (i.p.) infected with 5×10^7^ conidia or 4×10^6^ hyphae of *F. pedrosoi* (final volume of 200 μl). Animals infected with 1x PBS were used as a control. After 3 hours, the animals were euthanized and the i.p lavage was performed with 5 mL of 0,05% PBS-FBS with 2 mM EDTA to prevent clots. The cells were spun down and the supernatant was collected to measurement the neutrophils chemoattractants keratinocyte-derived cytokine (KC) and macrophage inflammatory protein-1α (MIP-1α). The cells were washed and resuspended in 1x PBS and subjected to Neubauer’s chamber counting, followed by flow cytometry staining using anti-CD45, anti-CD11b and anti-Ly6G antibodies. To check whether TLR-2 and TLR-4 on neutrophils were directly responsible for neutrophils migration, we performed an in vitro assay using the transwell methodology. Briefly, conidia or hyphae were seeded into the bottom of a 24 well-plate and neutrophils were added to the transwell insert, which was then placed in the top of the well. After a three hour incubation, neutrophil migration was quantified according to manufacturer’s instructions (Costar 3472)

### In-vivo infection

To confirm that TLR-2 and TLR-4 are two important receptors in CBM infection, we i.p. infected WT, TLR-2KO and TLR-4KO animals with 5×10^7^ *F. pedrosoi* conidia of. After 24 hours, the animals were euthanized, and the spleen and liver were collected to analyze the cell populations and fungal load. Briefly, the organs were harvested and smashed through a cell strainer (70 μM). An aliquot of the organ-macerate was collected and seeded onto Sabouraud agar for further CFU analysis. The rest of the organ macerate was centrifuged and placed over a 3 mL of Ficoll (1119 density) to isolate the leukocytes from the other tissue cells. The leukocytes were collected from the top of the Ficoll layer and stained with the following antibodies: CD45^+^, CD11b^+^ and F4/80^+^ (macrophages); CD45^+^, CD11c^+^, MHCII^+^ and F4/80^-^ (dendritic cells); and CD45^+^, CD11b^+^ and Ly-6G^+^ (neutrophils).

## Results

### Neutrophil fungicidal activity against *F. pedrosoi* conidia and hyphae

To determine whether neutrophils were capable of killing *F. pedrosoi* conidia and hyphae, we incubated the fungal particles with (or without) WT purified neutrophils for 2 hours. After, the cells were lysed in distilled water and seeded onto Sabouraud agar for 5 days for CFU counting. Conidia and hyphae incubated without neutrophils were used to set the 100% CFU (100% survival, or 0% killing). The CFU counts showed that in two hours purified neutrophils were able to kill both conidia and hyphae (**Fig. 1 – top left panel**). We next questioned whether this conidial and hyphal fungicidal activity was TLR-2 and TLR-4-dependent. To answer that question, we repeated this experiment using WT, TLR-2KO and TLR-4KO neutrophils. Our data showed that the neutrophil killing activity of conidia was impaired in TLR-2KO and TLR-4KO cells (**Fig. 1 – top middle panel**), while hyphal killing was not impacted (**Fig. 1 – top right panel**). These results suggest that *F. pedrosoi* particles activates neutrophils by distinct pathways.

**Figure 1.**
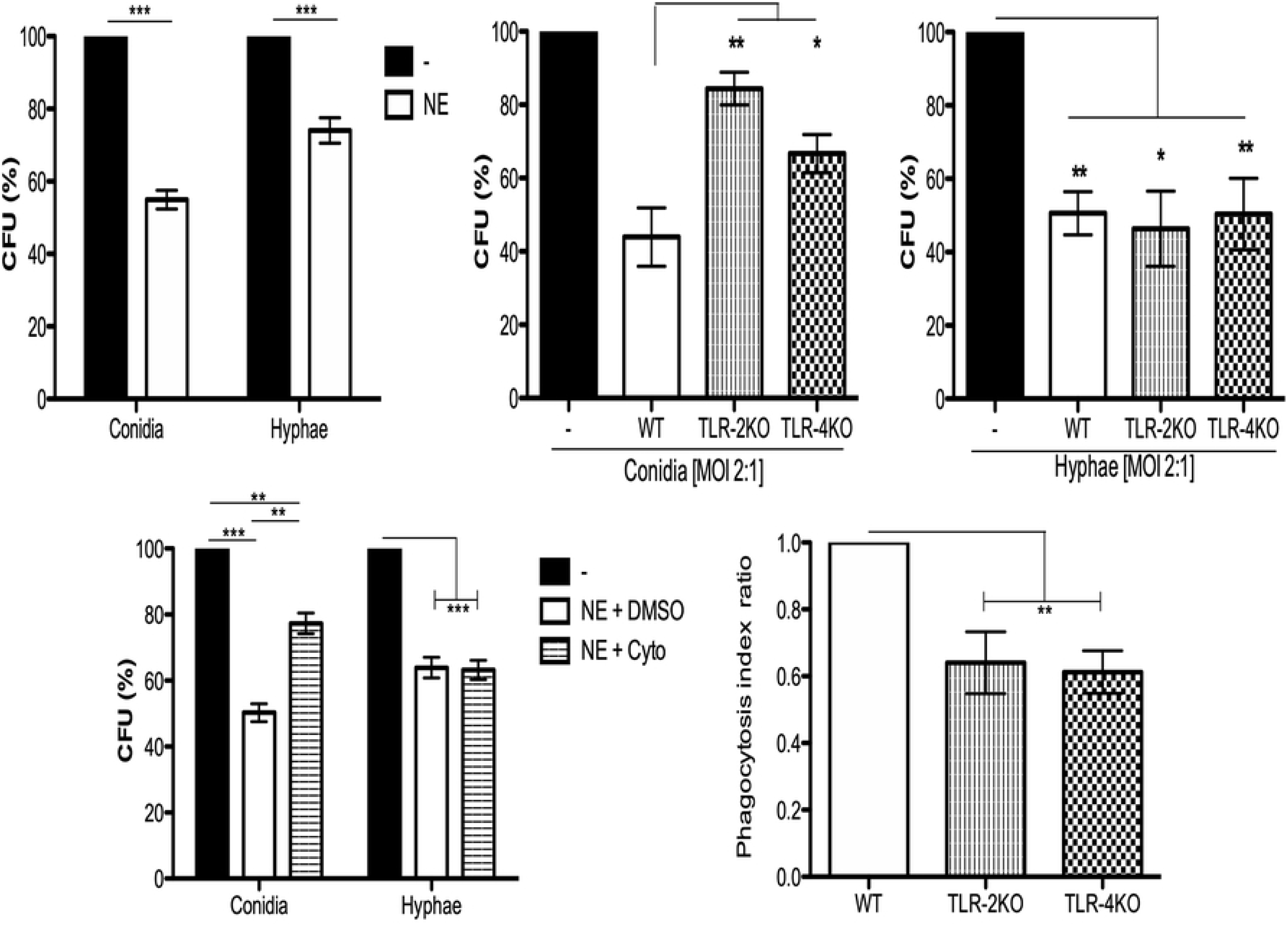
Neutrophils Toll like receptors 2 and 4 are important to kill conidia but not hyphae. (Top left panel) Neutrophils from WT were purified and incubated with conidia (MOI 1:2) or hyphae (MOI 1:1) of *F. pedrosoi* for 2 hours at 37° C. After, cells were lysed with sterile distilled water and seeded onto Sabouraud agar. After seed, plates were kept at 37° C incubators for 5 days prior to the final colony counting. As control, conidia and hyphae were incubated in the same conditions without neutrophils. Conidia and hyphae CFU obtained from controls were used as 100%. Using neutrophils from TLR-2KO and TLR-4KO animals we verified that these receptors are important against conidia (top middle panel) but not hyphae (top right panel) killing (left panel). Pre-incubating WT neutrophils with cytochalasin D (or DMSO – vehicle) for 30, we verified that phagocytosis is essential to kill *F. pedrosoi* conidia but not hyphae (bottom left panel). The importance of TLR-2 and TLR-4 in conidial phagocytosis was verified by immunofluorescence. Neutrophils were seeded over a coverslip and infected with conidia (MOI 2:1) for 2 hrs. After, the cells were fixed and permeabilized and the nuclei were stained with Sytox Green. The slides were mounted with Vecta-Shield ® and sealed with nail polish. At least 100 cells were analyzed to calculate the phagocytosis. Phagocytosis index observed in WT animals were set as 1 (100%) and the phagocytosis observed in TLR-2KO and TLR-4KO animals were compared to WT phagocytic index. (top left panel) n = 5, two-way ANOVA with Bonferroni’s post-test. \*\*\**p* < 0.001. n = 9, one-way ANOVA with Bonferroni’s post-test. \**p* < 0.05; \*\**p* < 0.01; \*\*\**p* < 0.001.

### Phagocytosis is a TLR-2 and TLR-4-depedent mechanism and is important for conidial but not hyphal killing

Our first hypothesis was that conidia were being killed via phagocytosis, while hyphae were not able to be internalized because of their size. Therefore, we performed a killing assay using cytochalasin D, a drug well-known to inhibit actin and myosin polymerization, thus inhibiting the phagocytosis process. Purified WT neutrophils were first incubated with cytochalasin D (or DMSO as a vehicle control) for 15 minutes, and then incubated for 2 hours with (or without) conidia or hyphae. Conidia and hyphae incubated without neutrophils were used to set the CFU as 100% (100% of survival). We demonstrated that the phagocytosis process was responsible, at least in part, for conidial but not hyphal killing (**Fig. 1 – bottom left panel**). To check whether the TLR-2 and TLR-4 were important for the phagocytic activity, purified neutrophils from WT, TLR-2KO and TLR-4KO animals were used to perform an immunofluorescence assay to quantify the phagocytic index of these cells. First neutrophils were seeded onto a coverslip and incubated for two hours with conidia (MOI 1:4). After cells were fixed and permeabilizated, the nuclei were stained with Sytox Green and the phagocytosis index was analyzed by counting 100 cells per group. The phagocytosis index obtained for WT neutrophils was set to a ratio of 1 (or the 100 % phagocytosis index). The TLR-2KO and TLR-4KO phagocytosis indexes were calculated and then compared to the WT index. Our results showed that conidia phagocytosis is impaired to approximately 35% to 45% in TLR-2KO and TLR-4KO neutrophils compared to WT neutrophils (**Fig. 1 – bottom right panel and Suppl. Fig. 1)**.

### *F. pedrosoi* conidia stimulate, while hyphae block neutrophils ROS production

ROS production is a well-described mechanism by which neutrophils and others phagocyte uses to kill different types of pathogens. Although ROS production is usually associated with the phagocytosis process, it is known that phagocytes can also release ROS extracellularly, killing unphagocytosed pathogens. Thus, we performed luminol-enhanced chemiluminescence assays to verify whether neutrophils were producing ROS during infections with conidia and hyphae. First, we seeded the neutrophils in the presence of luminol reagent, and then the cells were then stimulated with medium (negative control), conidia (MOI 1:2) or hyphae (MOI 5:1). After 90 minutes, the area under the curve was used to calculated the total ROS production. Our data showed that conidia stimulated neutrophils ROS production, while hyphae did not (**Fig. 2 - top left panel**). In fact, hyphae seem to block ROS production, leading to a level of ROS that was lower than the unstimulated cells. To confirm if hyphae were acting to block ROS production, we stimulated neutrophils with PMA, an agonist of NADPH-oxidase. Neutrophils stimulated with PMA showed a high level of ROS production; however, neutrophils stimulated with PMA in the presence of hyphae showed a statistically lower level of ROS production, confirming that hyphae were acting to block ROS production even in the PMA-stimulated cells (**Fig. 2 – top right panel**). Repeating these experiments using heat-killed hyphae, we demonstrated that only live hyphae have the capacity to block ROS production (**Fig. 2 – bottom**).

**Figure 2.**
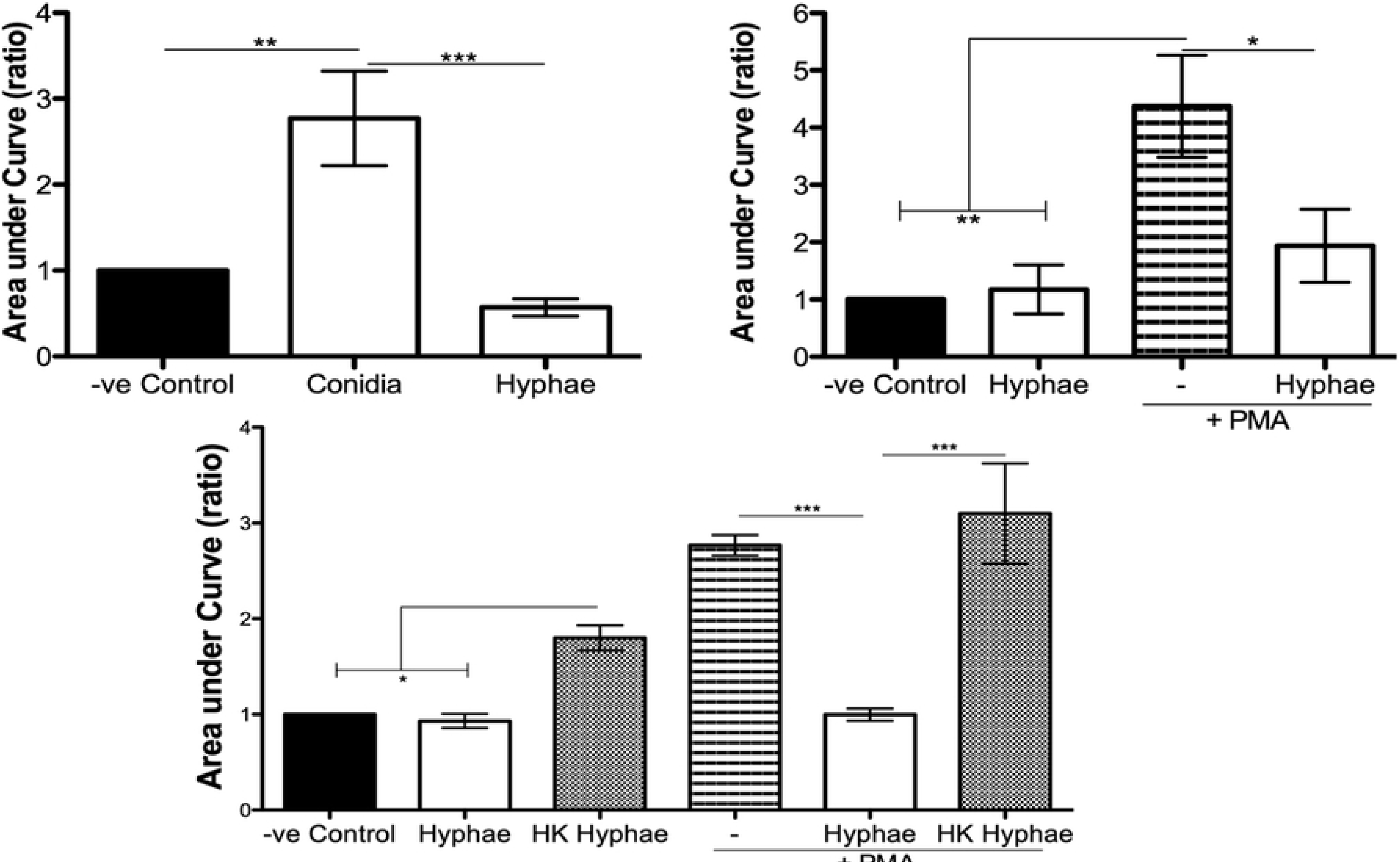
Neutrophil ROS production is stimulated by *F. pedrosoi* conidia and blocked by *F. pedrosoi* hyphae. (Top left panel) Purified WT neutrophils were seed in a 96 well-plate in the presence of luminol reagent and stimulated with *F. pedrosoi* conidia or hyphae. The ROS production was measured every 2 minutes to 60 minutes. As unstimulated control, neutrophils were incubated in the absence of fungi, to measure the ROS production during the steady-state. The area under the curve was calculated to measure the total ROS production after 60 minutes. (Top right panel) To confirm that hyphae block ROS production, we stimulated the cells with PMA (highly-stimulated ROS production) in the presence or absence of hyphae. (Bottom panel) Using heated-killed (HK) hyphae we demonstrated that live hyphae blocks, while HK hyphae stimulates, ROS production. n = 3, one-way ANOVA with Bonferroni’s post-test. \**p* < 0.05; \*\**p* < 0.01 \*\*\**p* < 0.001.

### ROS production during conidial infection is TLR-2 and TLR-4-dependent

Because conidia phagocytosis and killing was impaired in TLR-2KO and TLR-4KO neutrophils, we checked whether the ROS production was affected in these cells. Using the luminol-enhanced chemiluminescence assay, we verified that ROS production was impaired in TLR-2KO and TLR-4KO neutrophils infected with *F. pedrosoi* conidia of (**Fig. 3 – top left panel**). We also verified that the capacity of hyphae to block neutrophil ROS production occurred via a mechanism independent of TLR-2 and TLR-4 (**Fig. 3 – top right panel**).

**Figure 3.**
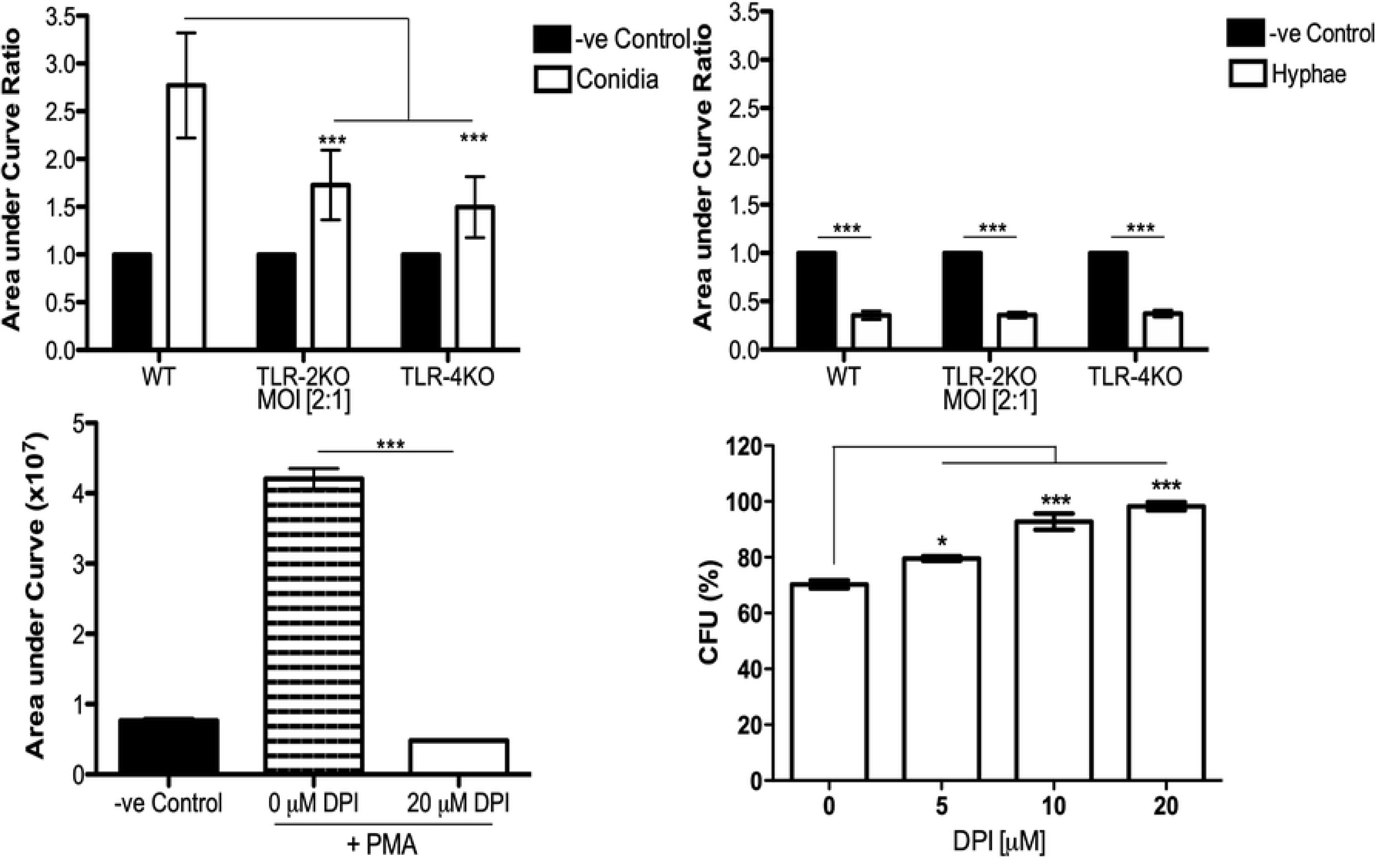
Neutrophil ROS production by *F. pedrosoi* conidia is a mechanism relied on TLR-2 and TLR-4 and essential to conidial killing. Purified WT, TLR-2KO and TLR-4KO neutrophils were seed into a 96 well-plate in the presence of luminol reagent and stimulated with *F. pedrosoi* conidia (top left panel) or hyphae (top right panel). As unstimulated control, neutrophils were incubated in the absence of fungi, to measure the ROS production during the steady-state. The ROS production was measured every 2 minutes to 60 minutes and the area under the curve was calculated to measure the total ROS production after stimulation. The area under the curve ratio was calculating using the control of each group (WT, TLR-2KO or TLR-4KO) and setting as 1. n = 5, two-way ANOVA with Bonferroni’s post-test. \*\*\**p* < 0.001. To verified whether conidial killing was dependent on ROS production, we used a range of concentrations of DPI a potent NADPH-oxidase inhibitor. First, we incubated WT neutrophils with DPI (or not) and stimulated them with PMA. The ROS production was measured to confirm the inhibition activity of the drug (bottom left). Then, using a range of DPI concentration (0μM - 20μM) we analyze the importance of ROS production in conidial killing. After pre-incubation with DPI, the purified neutrophils were incubated with conidia for 2 hrs. After incubation, the cells were lysed with distilled water and the supernatant was seeded onto Sabouraud agar at 37° C for 5 days (bottom right). n = 4, one-way ANOVA with Bonferroni’s post-test. \**p* < 0.05; \*\*\**p* < 0.001.

### Conidia killing is ROS-dependent

To confirm that conidia killing was occurring due to ROS production, we performed a killing assay using different concentrations of DPI, a NADPH-oxidase inhibitor drug. First, we performed an ROS assay in the presence (or absence) of 20 μM DPI. Neutrophils were previously incubated with medium (negative control) and 0 μM (DMSO as a vehicle) or 20 μM of DPI for 15 minutes. After, the cells were stimulated with PMA to confirm that this DPI concentration was able to completely block ROS production (**Fig. 3 – bottom left panel**). Next, the killing assay was performed using DPI at concentration ranging from 0 μM – 20 μM of DPI. Our data shows that in the absence of ROS production, the conidial survival was approximately 90% – 95%, proving that ROS is essential for conidia killing (**Fig. 3 – bottom right panel**).

### NETs release during *F. pedrosoi* hyphae infection is TLR-2 and TLR-4-independent

Although phagocytosis and ROS production were responsible for conidial killing, these mechanisms were not involved in hyphal killing. Thus, we asked whether hyphae were stimulating NETs release. Thus, a DNA release assay was performed using Sytox Green to quantify NETs release during infection with conidia (MOI 1:2) and hyphae (MOI 5:1). We first verified that hyphae, but not conidia were inducing NETs release (**Fig. 4 – top left panel**). Then, using TLR-2KO and TLR-4KO neutrophils we showed that these receptors are not responsible for neutrophil activation and NETs release (**Fig. 4 – top middle and right panel**). NETs release by WT neutrophils was also verified by immunofluorescence microscopy (**Fig. 4 middle**).

**Figure 4.**
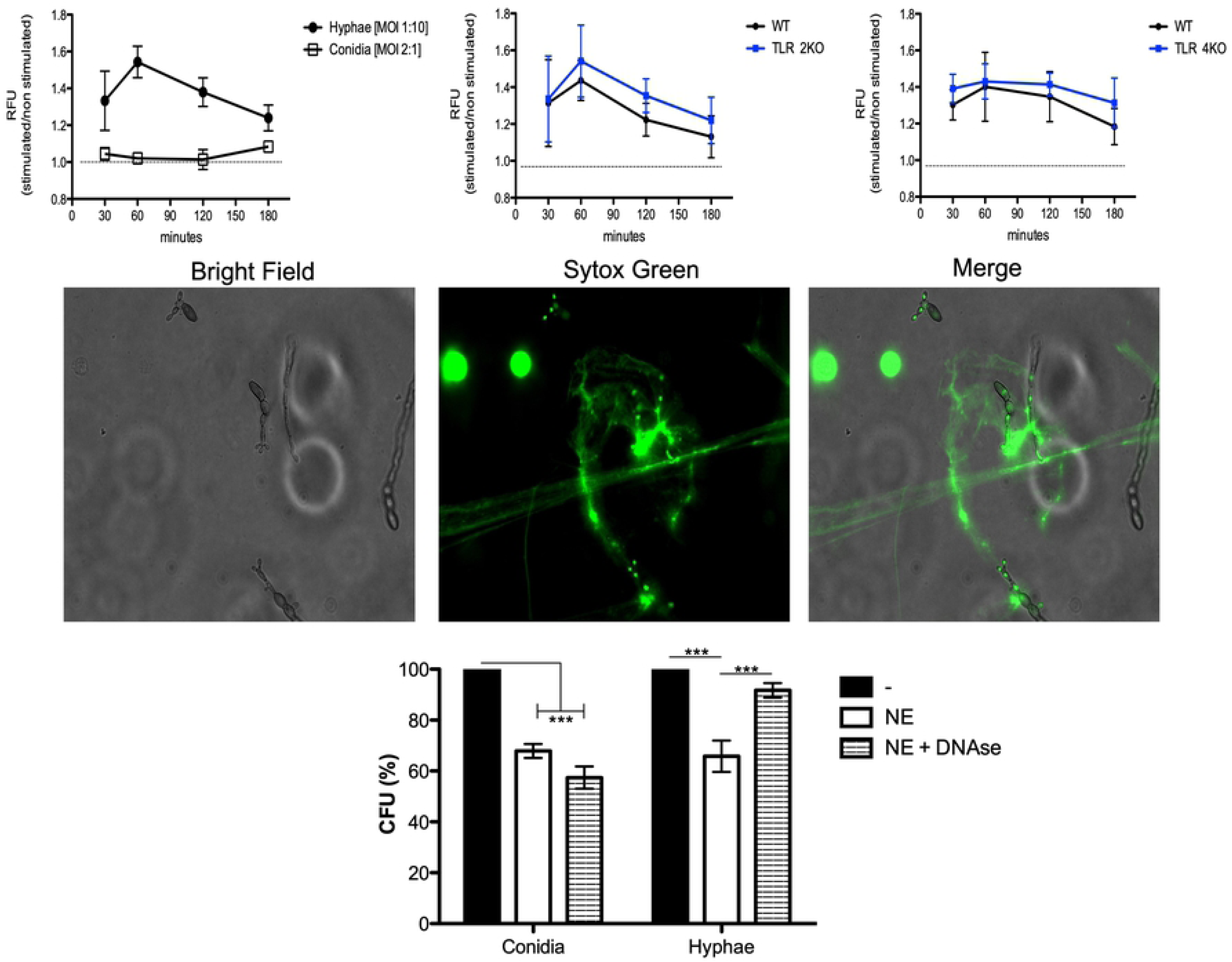
NETs release against *F. pedrosoi* hyphae infection is a mechanism independent of TLR-2 and TLR-4 and responsible for hyphae killing. (Top panels) Purified WT neutrophils were resuspended in media containing 5 µM Styox Green dye in resting condition (dashed lines - negative control) or incubated with *F. pedrosoi* hyphae or conidia. Florescence was recorded by a plate reader for every 30 min up to 3 h. Ratio of DNA release (NETotic index) shows NETosis over hyphae but not conidia infection (top left panel). After 180 min, the neutrophils incubated with hyphae were fixed with 4% (v/v) PFA for 15 min and analyzed by immunofluorescence microscopy (middle panel). Using TLR-2KO and TLR-4KO neutrophils we verified that NETs release is a mechanism independent from TLR-2KO and TLR-4KO (top middle and right panel). To confirm that NETs kills *F. pedrosoi* hyphae (but nor conidia) WT purified neutrophils were incubated with *F. pedrosoi* conidia or hyphae, in the presence or absence of DNase (bottom panel). After 2 hrs the cells were lysed with distilled water and seeded in Sabouraud agar at 37° C for 5 days. Conidia and hyphae incubated only with media (no neutrophils) was set as 100% CFU. n = 5, two-way ANOVA with Bonferroni’s post-test. \*\*\**p* < 0.001.

### NET released by neutrophils kills *F. pedrosoi* hyphae

Although neutrophils are able to release NETs against several pathogens, it has been shown that some pathogens are able to degrade or evade killing by NETs release. Therefore, to show that the NETs released in response to *F. pedrosoi* hyphae have fungicidal activity and can kill the fungal particles, we performed a killing assay in the presence of DNase. As expected, the survival index of conidia did not change, considering that we showed that conidia did not stimulated NETs release (**Fig. 4 bottom**). However, a statistical increase in hyphal survival was demonstrated when we disrupted NETs with DNase, confirming that NET released by neutrophils has fungicidal activity against *F. pedrosoi* hyphae.

### Neutrophils’ migration is impaired in TLR-2KO and TLR-4KO animals infected with *F. pedrosoi*

Neutrophils are known to be the first cell to migrate to the infection site. To tested whether TLR-2 and TLR-4 play a role in neutrophil migration we i.p infected WT, TLR-2KO and TLR-4KO animals with conidia and hyphae for 3 hours. After, we recovered the migrated cells by i.p. lavage, and the cells were counted and stained for flow cytometry analysis. Our results showed a higher neutrophil influx during infection with hyphae, compared to conidia. A severe impairment in neutrophils migration was observed in animals lacking TLR-2 and TLR-4 (**Fig. 5 – top panels**). To check whether TLR-2 and TLR-4 directly affect neutrophil migrations, we performed an in vitro migration assay using the transwell methodology. Briefly, conidia or hyphae were seeded into the bottom of a 24 well-plate and neutrophils were added to the upper compartment (inside the insert) that was placed in the top of the well. After three hours of incubation, no neutrophils migration towards the fungus was observed, suggesting that conidia and hyphae cannot stimulate neutrophils migration *per se* in an in vitro assay (data not shown). To analyze whether TLR-2 and TLR-4 indirectly stimulate neutrophils migration towards the site of conidial and hyphal infection, we measured the levels of KC (**Fig. 5 – middle panels**) and MIP-1α **(Fig. 5 – bottom panels)** in the peritoneal lavage fluid after 3 hours of infection. Our data suggests that TLR-2 and TLR-4 are important receptors for sensing *F. pedrosoi* and thus, they stimulate the production of chemokines such as KC and MIP-1α by resident cells present in the peritoneum. These impaired in KC and MIP-1α production by TLR-2KO and TLR-4KO animals affected neutrophils migration to the infection site.

**Figure 5.**
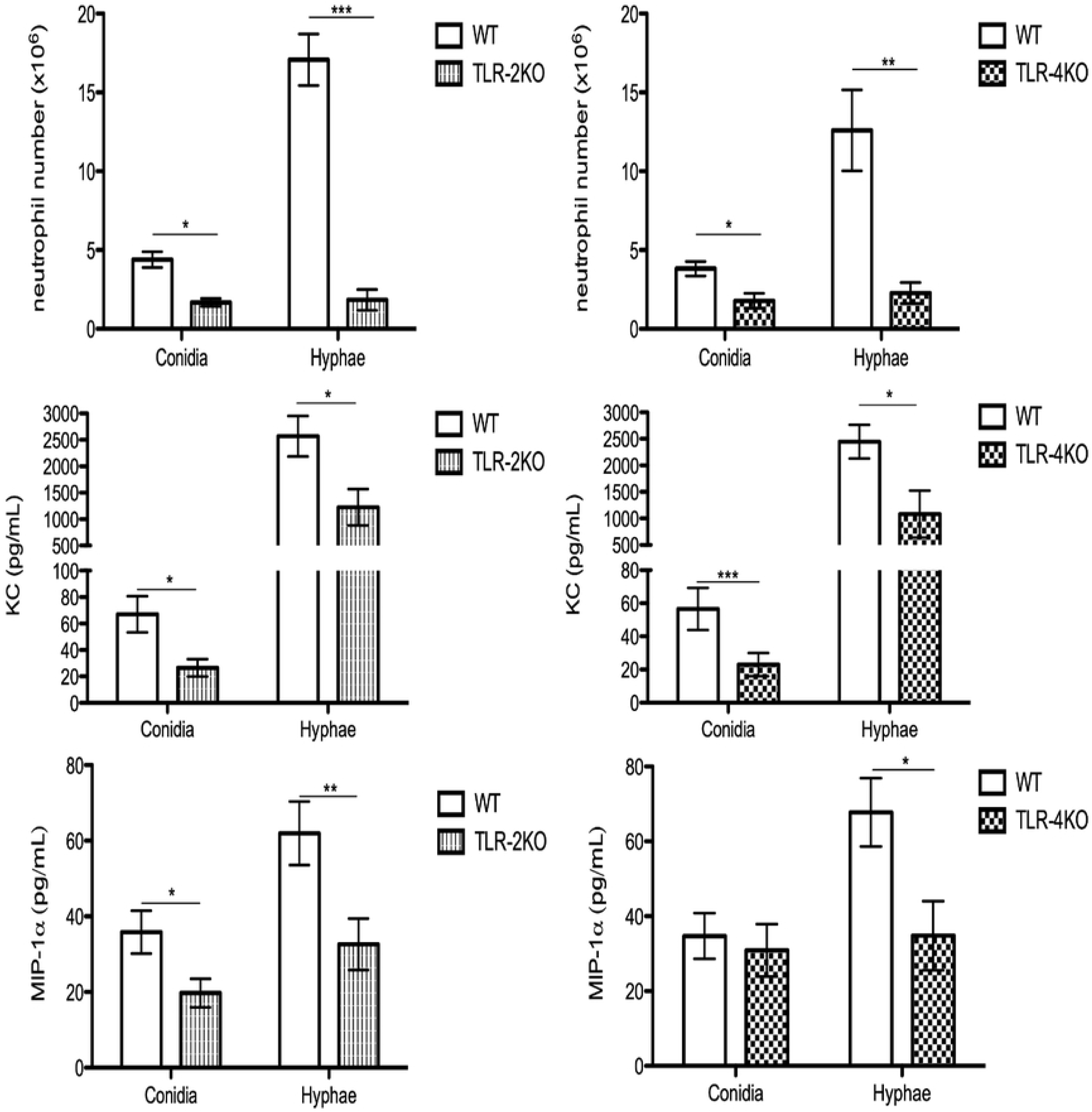
Neutrophil attraction to conidia and hyphae infected site is dependent on TLR-2 and TLR-4 and chemokines KC and MIP-1α. WT, TLR-2KO and TLR-4KO animals were i.p. infected with *F. pedrosoi* conidia or hyphae. After 3 hours of infection, animals were euthanized and the i.p. lavage were collected with PBS 5% FBS 2mM EDTA. The lavage was centrifuged and supernatant was collected to measurement of chemokines by ELISA (middle and bottom panels) and the cells were harvested and then counted in Neubauer chamber and stained with anti-CD45, anti-ly6G and anti-CD11b to detect neutrophils migration by flow cytometry technique (top panel). n = 9-13, two-way ANOVA with Bonferroni’s post-test. \**p* < 0.05; \*\**p* < 0.01; \*\*\**p* < 0.001; t-student. \**p* < 0.05; \*\**p* < 0.01; \*\*\**p* < 0.001.

### Higher fungal burden in spleen and liver of TLR-2KO and TLR-4KO infected animals

To confirm that TLR-2 and TLR-4 are important in controlling CBM infection in vivo, we i.p. infected WT, TLR-2KO and TLR-4KO animals with *F. pedrosoi* conidia for 24 hours. After, the spleen and liver were harvested, and an aliquot was seeded onto Sabouraud agar plates for later CFU counting. Our results showed that animals lacking TLR-2 and TLR-4 had higher fungal loads in the spleen and liver compared to WT animals, confirming that these receptors were also important for controlling the disease in murine models (**Fig. 6**). An increase in the neutrophil population was seen in the spleens and livers of the KO animals (**suppl. Fig. 2**).

**Figure 6.**
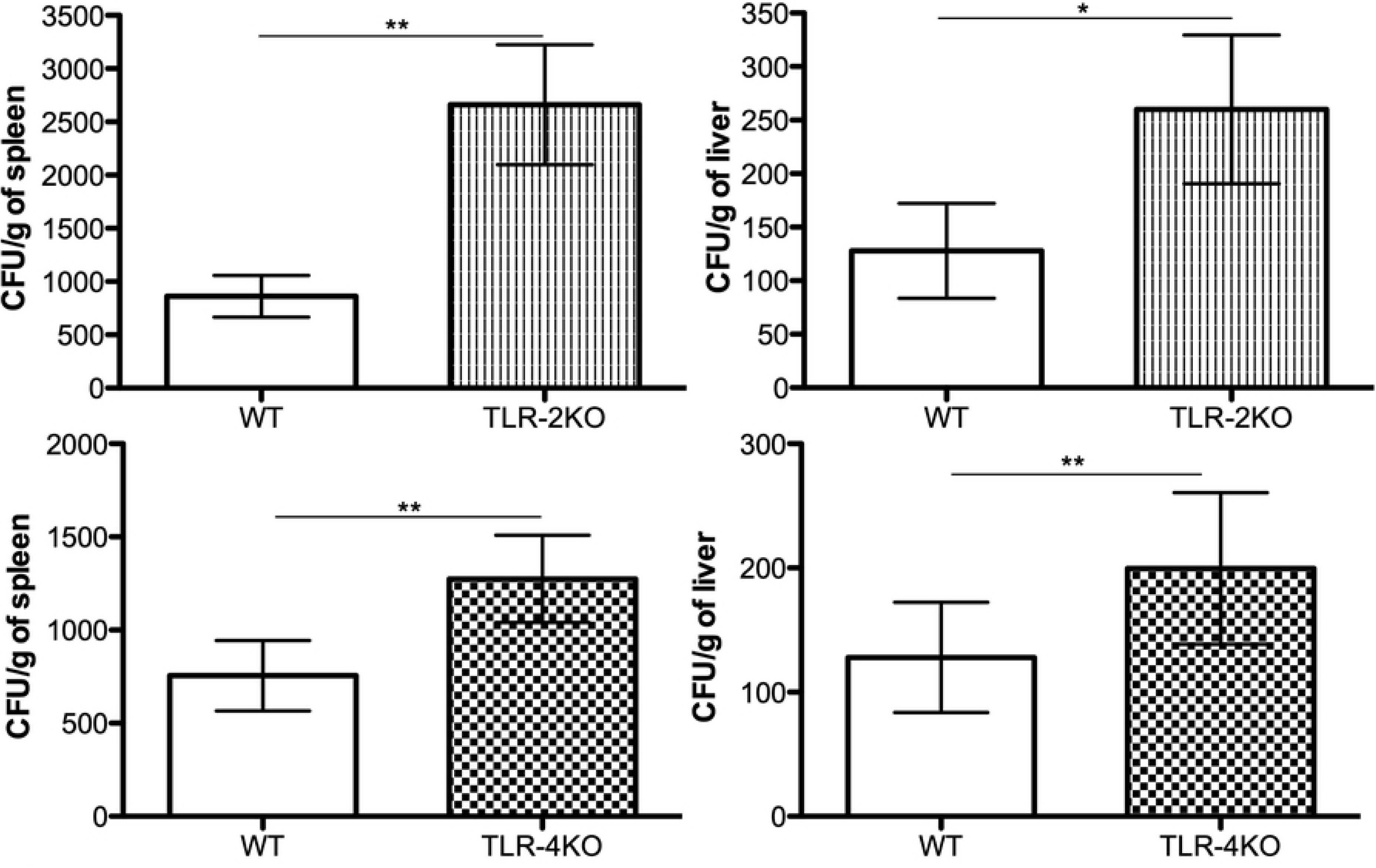
TLR-2 and TLR-4 are essential for fungal load control in early *F. pedrosoi* conidia infection. WT, TLR-2KO and TLR-4KO animals were infected i.p. with 5×10^7^ *F. pedrosoi* conidia. After 24 hours, the animals were euthanized and the liver and spleen were harvest. An aliquot was seeded onto Sabouraud agar to analyze the fungal load in these animals. n = 10, t-student. \**p* < 0.05; \*\**p* < 0.01.

## Discussion

Currently, CBM treatment has low cure rates and is based on multidrug prescriptions and, in some cases, cryo/heat therapy with surgery (10–14). More effective treatment is needed, therefore, a better understanding of the host-pathogen interaction is crucial. Although several studies have demonstrated the essential roles of TLR-2, TLR-4 (35–37) and neutrophils in several infections (38), including fungal infections (20), the real significance of these receptors and cell in CBM disease has not been fully addressed to date. In the present study, we elucidated the roles of TLR-2 and TLR-4 in several neutrophil effector functions, and described for the first time the mechanism used by neutrophils to kill *F. pedrosoi* hyphae.

The first study of neutrophils function in *F. pedrosoi* infection was carried out in 1996 (34). In that work, the authors demonstrated that conidia were killed by neutrophils through the production of extracellular ROS once a few particles had been detected inside neutrophils. Our study confirmed the fungicidal activity of neutrophils towards conidial infection and demonstrated for the first time that neutrophils also have fungicidal activity against *F. pedrosoi* hyphae (**Fig. 1**). Our results also confirmed that ROS production is essential for conidia killing (**Fig. 3**). However, different from previous studies, we showed a high neutrophil phagocytic activity (**Fig. 1**) and demonstrated that this process is essential for eliminating conidia particles (**Fig. 1**). Although some of our results disagree with previously published data, we have to consider that Rozental and colleagues used rat neutrophils and some results may be species-specific.

Although the vast majority of studies focus on understanding the roles of TLR-2 and TLR-4 in bacterial infections (once that these receptors are known to recognize peptidoglycan and lipopolysaccharide, respectively), these receptors are also important in fungal infections, because they bind to glucan/mannan and rhamnose, respectively (39, 40). Our findings showed that TLR-2 and TLR-4 were essential to conidial but not hyphal killing (**Fig. 1**). These receptors were also found to be important for killing *Aspergillus fumigattus* conidia (41) and *Candida albicans* blastocinidia (42), however, *C. albicans* hyphae are recognized by only TLR-2, while *A. fumigatus* hyphae are recognized by only TLR-4 (42). Its is known that different forms of the same fungal species may be presented in the environment or in the host during an infection process (such as conidia/hyphae or yeast/blastoconidia/pseudohyphae). In additional to their difference in size, the cell-wall components of these differet fungal forms can be very different (43), and this seems to be crucial to fungal recognition by the host-immune system. Although proteomics studies of the *F. pedrosoi* cell wall have been relatively rare, a couple of studies have demonstrated that conidia and hyphae from *F. pedrosoi* present different cell wall compositions, with some similarity in the components but different levels of expression (44, 45). Therefore, although the TLR-2 and TLR-4 are crucial in conidial killing, probably due to the difference in cell wall components, these receptors does not play a role in *F pedrosoi* hyphal killing (**Fig. 1**). Using immunofluorescence microscopy we verified that the absence of TLR-2 and TLR-4 leads to impaired conidial phagocytosis (**Fig. 1 and Suppl. Fig. 1**). Similar results were seen in *P. brasiliensis* (46, 47) and *Sporothrix brasiliensis* (48, 49) infection. By contrast, in *C. albicans* and *A. fumigatus* infection TLR-4 does not play a crucial role in the phagocytic process (50, 51).

Although phagocytosis is an important neutrophil effector function, it is not sufficient for particle killing. For that, the phagosome has to fuse with the lysossome so that the oxidative burst can take place. This leads to ROS and reactive nitrogen species (RNS) production, which is responsible for the killing of phagocytosed pathogens. However, it is well described that phagocytes can also be activated and release ROS and RNS extracellularly (phagocytosis-independently), making this a possible mechanism of extracellular hyphae killing (52). Based on that, we asked whether conidia and hyphae were stimulating neutrophil ROS production. Our findings showed that neutrophils produced ROS during conidial infection in a TLR-2 and TLR-4-dependent manner (**Fig. 3**). Similar data were published for *Sporothrix brasiliensis* infection, in which the authors showed that these receptors were important for the pathogen phagocytosis, with impaired RNS production observed in TLR-2KO and TLR-4KO macrophages (53, 54). Since these receptores were not important for *C. albicans* phagocytosis (51), ROS production was not affected by the absence of the TLR-2 and TLR-4 receptors (55). Our results demonstrated that althought neutrophils are able to release extracellularly ROS and RNS, *F. pedrosoi* hyphae do not stimulate neutrophil ROS production (**Fig. 2**). In fact, we observed that neutrophils infected with hyphae were producing lower amounts of ROS than resting neutrophils (unstimulated). Some studies have demonstrated that conidia have the capacity to block nitric oxide (NO) production even in a IFN-γ-stimulated murine macrophages (56, 57). Even though conidia and melanin purified from conidia were shown to block NO production, these particles were found to stimulate macrophage ROS production. However, the authors did not show the basal levels of ROS-production in healthy animals (uninfected), therefore, we can not conclude whether the hyphae were weakly, not stimulating or blocking ROS production (58). Thus, by preactivating bone marrow-purified neutrophils with PMA, we showed that hyphae block neutrophil ROS production. This inhibition is lost when the hyphae are heat-killed (**Fig. 2**). Similar results were found with *Aspergillus nidulans* hyphae, where no ROS was produced by infected neutrophils. The authors verified that *Aspergillus nidulans* hyphal killing by NADPH-oxidase deficient neutrophils (from patients with granulomatous chronic disease) was similar to healthy neutrophils, demonstrating that neutrophils kill *A. nidulans* hyphae in a ROS-independent manner (59). Our findings also show that TLR-2 and TLR-4 are not involved in the blocking of ROS production (**Fig. 3**). Therefore, our findings indicates that hyphae are killed by a ROS-independent mechanism. Even though the first description of NETs release has suggested that this activity was dependent on ROS production by NADPH-oxidase (60), several more recent studies demonstrated that NETs release can be an NADPH oxidase-dependente or -independent process (61–63). An important study demonstrated that neutrophils can sense the size of a pathogen to decide whether the cells will phagocytose or release NETs to kill the pathogen (64). However, pathogen size is not the only feature that leads to neutrophil activation and NET release, because some bacteria (60) and yeast (22) stimulate neutrophil NET-release even though they are small enough to be phagocytosed. Based on that, we asked whether *F. pedrosoi* conidia and hyphae of were stimulating NETs release. Our findings show that neutrophils release NETs in response to *F. pedrosoi* hyphae but not conidia. Unlikely phagocytosis, NETs release occurs via a TLR-2 and TLR-4 independent mechanism (**Fig. 4**). Although neutrophils realease NETs fibres during several pathogen infections, different studies have demonstrated that some pathogens can evade NETs killing, by degrading the fibres, through the formation of biofilms or as a result of the presence of extracellular capsule (65). Therefore, we performed a killing assay with DNase and demonstrated for the first time that NETs are a mechanism used by neutrophils to eliminate *F. pedrosoi* hyphae. We also performed in vivo experiments to evaluate the roles of TLR-2 and TLR-4 in a murine CBM infection model. First, we verified that there was impaired neutrophil migration to the infection site in animals lacking TLR-2 and TLR-4 in both conidial and hyphal infections (**Fig. 5**). We showed that the impared neutrophil migration was due to the decreased levels of KC and MIP-1α released by the resident peritoneal cells (**Fig. 5**). At least in our hands, conidia and hyphae could not directly stimulate neutrophil migration in in vitro transwell asssays (data not shown). Similar results were observed in *A. fumigatus* infection, where TLR-2KO and TLR-4KO macrophages realeased lower levels of the MIP-2 chemokine, resulting in a decrease in neutrophils migration (66). In *C. albicans* infection, animals lacking TLR-4 showed lower levels of MIP-2 and KC leading to impared neutrophil migration to the infection site (51). Finally, our study showed that TLR-2 and TLR-4 are important in controlling acute CBM infection, because animals lacking these receptors had higher spleen and liver fungal loads (**Fig. 6**).

In summary, our results show for the first time that neutrophils are important for *F. pedrosoi* conidia and hyphae killing. The cell wall composition and pathogen size may be acting to modulate neutrophil function, leading to phagocytosis and ROS production during conidial infection, while ROS-independent NETs release is the main efffector function involved in hyphal killing. We also demonstrated that TLR-2 and TLR-4 are important receptors in rcognition of conidia but not in recognition or killing of hyphae. Theses receptors were also crucial for neutrophil migration towards the infection site and in the control of the fungal burden in the animals. Therefore, our findings help to better understand the physiopathology of CBM and how the neuturophils fight against *F. pedrosoi* conidia and hyphae infection.

## Abbreviations

CBM: Chromoblastomycosis
CFU: colony-forming unit (CFU)
DPI: Diphenyleneiodonium
HK: heat-killed
i.p.: intraperitoneal
MOI: multiplicity of infection
NET: Neutrophil Extracellular Trap
PFA: Paraformaldehyde
ROS: reactive oxygen species

## Authorship

L.B. and S.A. designed the research, interpreted the data and wrote the manuscript. L.B., C.B., J.A., L. P., G.J., I.M. and R.A. performed the experiments. N.C. and K.F. provided scientific input. All authors read and approved the manuscript.

## Acknowledgments

This work was supported by the “Fundação de Amparo à Pesquisa do Estado de São Paulo” (FAPESP - process 2012/18598-7, 2014/11146-9, 2016/047293), “Coordenação de Aperfeiçoamento de Pessoal de Nível Superior” (CAPES) and “Conselho Nacional de Desenvolvimento Científico e Tecnológico” (CNPq).

**Figure Suppl 1** Conidia phagocytosis is a TLR-2 and TLR-4-dependent process. WT, TLR-2KO and TLR-4KO neutrophils were incubated with *F. pedrosoi* conidia (MOI 1:4) for 120 minutes. After incubation, the cells were washed and fixed with 4% PFA for 15 minutes followed by permeabilized with 0,01% PBS-T for 10 minutes. After washing, the neutrophils nuclei were stained with Sytox Green and the slides were mounted using Vecta-Shield ® and sealed with nail polish. At least 100 cells were analyzed to calculate the phagocytosis index showed in Fig. 1.

**Figure Suppl 2** The absence of TLR-2 or TLR-4 increase neutrophil population in vivo infection by conidia of *F. pedrosoi*. WT, TRL-2KO and TLR-4KO animals were infected i.p. with 5×10^7^ conidia of *F. pedrosoi*. After 24 hours, the animals were euthanized and the liver and spleen were harvest and the DCs, macrophages and neutrophil population was analyzed by flow cytometry. n = 10, two-way ANOVA with Bonferroni’s post-test. \**p* < 0.05; \*\**p* < 0.01; \*\*\**p* < 0.001.

